# Integration of Euclidean and path distances in hippocampal maps

**DOI:** 10.1101/2023.08.01.551513

**Authors:** L. Ottink, N. de Haas, C.F. Doeller

## Abstract

The hippocampus is a key region for forming mental maps of our environment. These maps represent spatial information such as distances between landmarks. A cognitive map can allow for flexible inference of spatial relationships that have never been directly experienced before. Previous work has shown that the human hippocampus encodes distances between locations, but it is unclear how Euclidean and path distances are distinguished. In this study, participants performed an object-location task in a virtual environment. We combined functional magnetic resonance imaging with representational similarity analysis to test how Euclidean and path distances are represented in the hippocampus. We observe that hippocampal neural pattern similarity for objects scales with Euclidean as well as path distance between object locations, suggesting that the hippocampus integrates both types of distances. One key characteristic of cognitive maps is their adaptive and flexible nature. We therefore subsequently modified path distances between objects using roadblocks in the environment. We found that hippocampal pattern similarity between objects adapted as a function of these changes in path distance, selectively in egocentric navigators but not in allocentric navigators Taken together, our study supports the idea that the hippocampus creates integrative and flexible cognitive maps.

## 1. Introduction

Navigating successfully in our everyday lives requires a mental representation of our environment. Such a cognitive map must contain information about relevant locations and relationships between them, in particular spatial distance. Furthermore, it must facilitate flexible behaviour, for instance planning detours when the environment changes^1–7^. In the current study, we firstly aim to investigate cognitive map formation by analysing neural representations of path as well as Euclidean distances. Secondly, we assess how these representations adapt when paths in the environment change.

Converging evidence shows that hippocampal activity correlates with the path distance to the goal during navigation^8–11^. The hippocampus also forms a map that stores distances between relevant locations^10–13^. Previous research^13^ suggested that the hippocampus forms a spatio-temporal distance map of object locations along a fixed path. Neural pattern similarity in the hippocampus increased the closer people remembered object-pairs to be^13^. Furthermore, it has also been found that hippocampal activity was modulated by Euclidean distance and estimated travel time between two landmarks^12^. However, both variables were highly correlated in this study, which makes it difficult to disentangle the effects. In the current study, we built on these previous studies, by asking participants to explore an environment freely instead of following a fixed path, and by separately manipulating Euclidean and path distances between objects. Both Euclidean and path distance are crucial aspects in real-world navigation. Path distances allow us to remember the shortest route, while Euclidean distances enable us to compute novel shortcuts. Here, we asked how Euclidean and path distances are simultaneously represented in the brain, and whether they are integrated. For this purpose, our participants performed a navigation task and learned eight locations in a virtual city^13, 14^.

Besides accurate representation of distances between relevant places, successful navigation requires a cognitive map that reacts flexibly to changes, for instance when a roadblock forces us to take a detour^1, 6^. The hippocampus signals the change in path distance during navigation^6, 10^. Here, our second aim was to test how hippocampal maps adapts when paths change. Therefore, participants performed a second navigation task in which we systematically altered path distances but not Euclidean distances between locations.

In both tasks, akin to Deuker and colleagues^13^, we used changes in neural pattern similarity between object locations as a proxy for the formation and update of map representations. We expected that lower distances would lead to higher similarity, and vice versa. Additionally, to assess cognitive maps behaviourally, participants performed two distance recall tasks and a location recall task.

Finally, we explored whether navigational strategies affect representations in the hippocampus. Therefore, we examined differences between response learners (egocentric navigators) and place learners (allocentric navigators)^4, 15–17^. These two strategies may result in different hippocampal signatures^17, 18^ based on the allocentric nature of Euclidean and egocentric nature of path distances. Map-like representations include Euclidean distances, involve place learning and relate to allocentric perspectives. Whereas route-like representations include path distances, involve response learning and relate to egocentric perspectives^15, 19, 20^. Therefore, hippocampal representations of response learners may be more sensitive to egocentric features (i.e., path distance), and those of place learners more to allocentric features (i.e., Euclidean distance). We adapted a T-maze task^16^ to categorise our participants into response and place learners, and tested for differential memory performance and neural representations of distance.

Taken together, we probed hippocampal representations of path and Euclidean distances, and how these might adapt after changes in the environment. Lastly, we explored whether different navigational strategies might yield different neural signatures in the hippocampus.

## 2. Methods

### 2.1 Participants

32 healthy participants (16 women) completed the experiments. Participants were recruited via the university’s online recruitment platform. Participants were between 18 and 32 years old (mean age = 22, SD = 3.1). At the beginning of the experiment, participants gave written informed consent to participate in the study and filled in a screening form to ensure that they did not meet any exclusion criteria for the MRI and behavioural labs. Participants were compensated at a rate of 8 Euro per hour of behavioural testing and 10 Euro per hour of MRI testing. The study was approved by the local ethics committee (CMO Regio Arnhem-Nijmegen, The Netherlands, nb. 2014/288).

All participants (13 place learners, 17 response learners and 2 mixed, as classified by the T-maze task) were included in the analyses of the behavioural tasks. Out of 32 participants, 30 participants (12 place learners, 16 response learners and 2 mixed) were included for analysing neural changes from Picture Viewing Task (PVT) 1 to PVT 2. 29 participants (12 place learners, 15 response learners and 2 mixed) were included for analysing neural changes from PVT 2 to PVT 3.

### 2.2 Experimental Design

We wanted participants to form a cognitive map of a large-scale virtual environment, including the locations of eight objects. To this end, we conducted an experiment that took place on two consecutive days (Figure 1A). During this experiment, participants completed in total three navigation tasks in a virtual city, where they had to learn 8 goal-locations and the shortest routes between them. The virtual city was an adapted (smaller) version of ‘Donderstown’ (Figure 1B)^13, 14, 21^. We programmed all navigation tasks with Unreal Development Kit (Unreal Engine 3, Epic Games, Inc.). The first navigation task, the training task, took place on day 1 (Figure 1A, C), and made sure the participants became familiar with the environment. During this task, participants navigated freely from goal-location to goal-location (presented as black boxes), in order to learn their positions and the shortest routes between them. The training task was followed by the Santa Barbara Sense of Direction Scale (SBSOD)^22^, a questionnaire to assess participant’s self-reported general navigational abilities and sense of direction. Day 1 ended with a T-Maze task to assess place and response learning (adapted from the version by Astur and colleagues^16^; Figure 1D). This allowed us to analyse whether there are differential behavioural and neural signatures of a cognitive map between users of the two navigational strategies (place versus response learning). On day 2, participants started with the second navigation task, the object location task (Figure 1A). This was similar to the Training task, with the same goal-locations, but now an object was presented when a participant arrived at the goal-location (Figure 1C, E). The participants thus learned the locations of eight objects. After the object-location task, participants performed a distance recall task, where they had to estimate Euclidean and path distance between each object-pair (Figure 1A). Subsequently, they performed the last navigation task, the so-called path change task. This was similar to the first object-location task, but here we manipulated the routes, and thus the path distances, between some object-pairs using roadblocks. At the end of day 2, participants performed a second distance recall task, and a bird-view placement task (Figure 1A). Here they had to place the eight objects on their remembered locations on a bird-view map of the virtual city. Before the training, and before and after the path change task, participants additionally performed a picture viewing task (PVT; Figure 1A). Here, the eight objects were repeatedly presented during an fMRI session. This allowed us to analyse the neural signatures of a cognitive map of the virtual environment. All tasks and analyses are described in more detail below.

**Figure 1.**
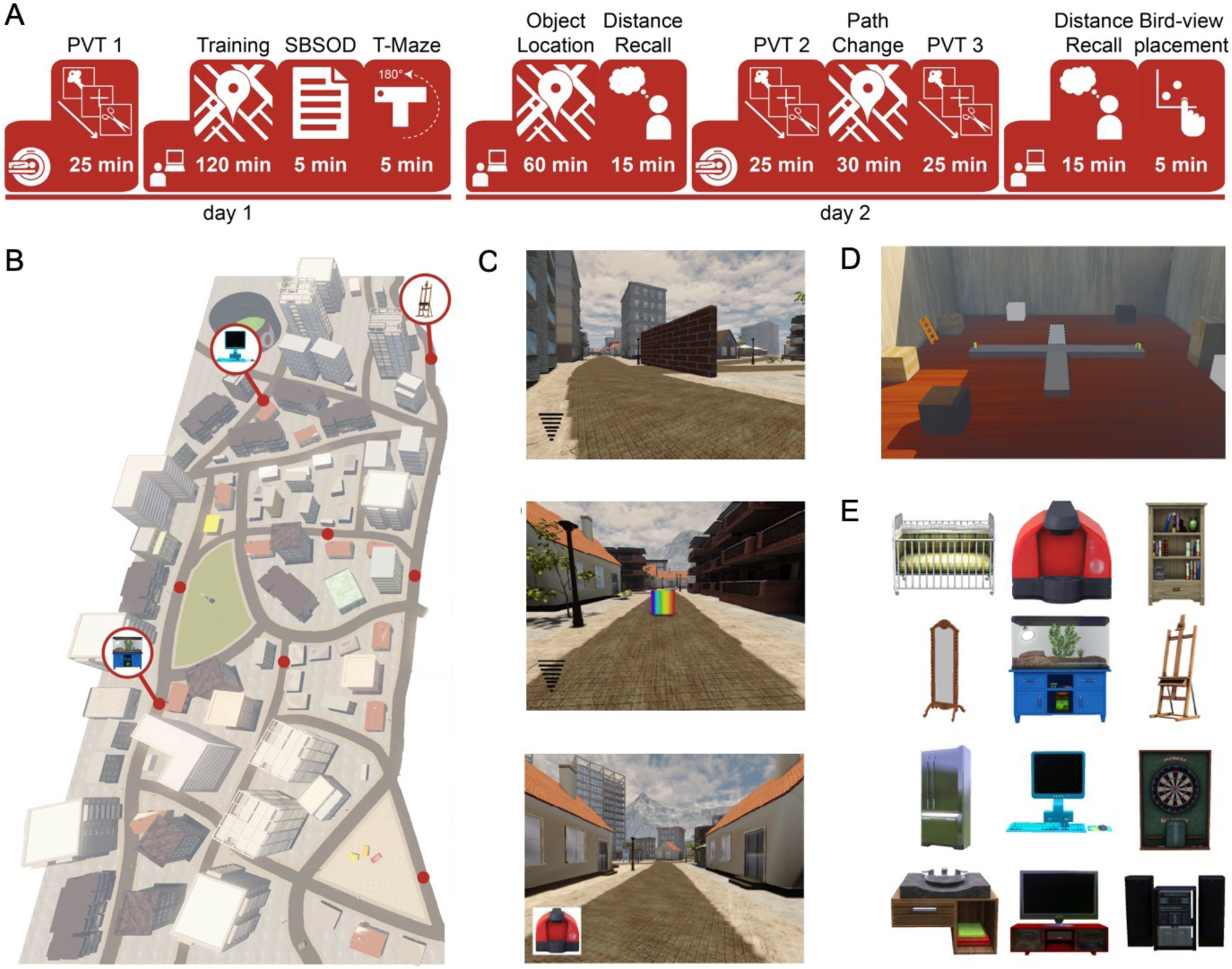
Experimental sessions and VR environment. A. Overview of the experimental sessions on day 1 and day 2. Day 1 started with a picture viewing task (PVT) during and fMRI session, followed by a behavioural session including the training task, Santa Barbara Sense of Direction scale and a T-maze task. On day 2, participants performed the object location task, followed by a distance recall task. The second PVT, path change task and third PVT were performed in the MRI-scanner. At last, participants performed a second distance recall and a bird-view task. B. Birdview of the virtual city. Object-locations are indicated with a red dot. Three example objects are shown. C. Screenshots of the navigation task. Top: navigation phase of the training task and first half of object-location task. The wifi-like signal indicates how close the participant is to the goal-location. The brick wall is one of the roadblocks. Middle: the box of the location would turn from black to multicoloured if it is the goal-location of the current trial. Bottom: Navigation phase during the second half of the object-location task, and the path change task. Instead of the wifi-like signal, participants are cued by the object of the goal-location. D. Screenshot of the T-maze task environment. During normal trials, the dark gray elevated platform was the T-maze, with a multicoloured box at the left and right arm. During probe trials, the T-maze was rotated 180°, and the light grey arm became the starting arm of the T. E. Eight out of twelve objects were randomly chosen for each participant. Each object was associated with one of the eight goal-locations in the virtual environment. The objects were: a baby bed, a coffee maker, a bookshelf, a mirror, a terrarium, a canvas stand, a fridge, a computer, a dart board, a sink, a TV and a stereo.

In our study, we wanted to assess whether participants formed a cognitive map of the virtual city. We did this by investigating neural representations of Euclidean and path distances between object-pairs, as both types of distances are important features of cognitive maps. To this end, we compared the representation of object-pairs that have a high versus low Euclidean distances, and of object-pairs that have a high versus low path distance. Apart from distinct Euclidean and path distances, the two distance types could also be integrated. It could be that the hippocampus does not distinctly represent Euclidean and path distances, but that it represents an integration of the two. It could thereby differentiate the extremes, and separate object pairs that have a high Euclidean as well as path distance from object pairs that have a low Euclidean as well as path distance. Since we manipulated some paths using roadblocks, some object pairs have relatively high Euclidean but relatively low path distance, or vice versa. Therefore, we furthermore compared object-pairs that have a high Euclidean and high path distance with object-pairs that have a low Euclidean and low path distance to test whether the two distance types are integrated. Hence, the ‘integration distance’ was the product of the Euclidean and path distance. 10 out of 28 object-pairs had the highest integration distance (i.e., a high Euclidean as well as path distance), and 11 out of 28 object-pairs had the lowest integration distance (i.e., a low Euclidean as well as path distance).

We experimentally disentangled Euclidean and path distances by placing three roadblocks in the virtual city (Figure 2A). Consequently, for some object-pairs, participants had to take a detour, while for others they could take the shortest route. In the path change task, we changed two of the three roadblocks. Now, participants could take the shortest route between some object-pairs where they previously had to take a detour, and vice versa (Figure 2A). These modifications allowed us to shorten the path distance for some object-pairs, and make the path distance longer for some other object-pairs in the path change compared to the training and object-location tasks. The Euclidean distance between the object-pairs remained the same for all navigation tasks.

**Figure 2.**
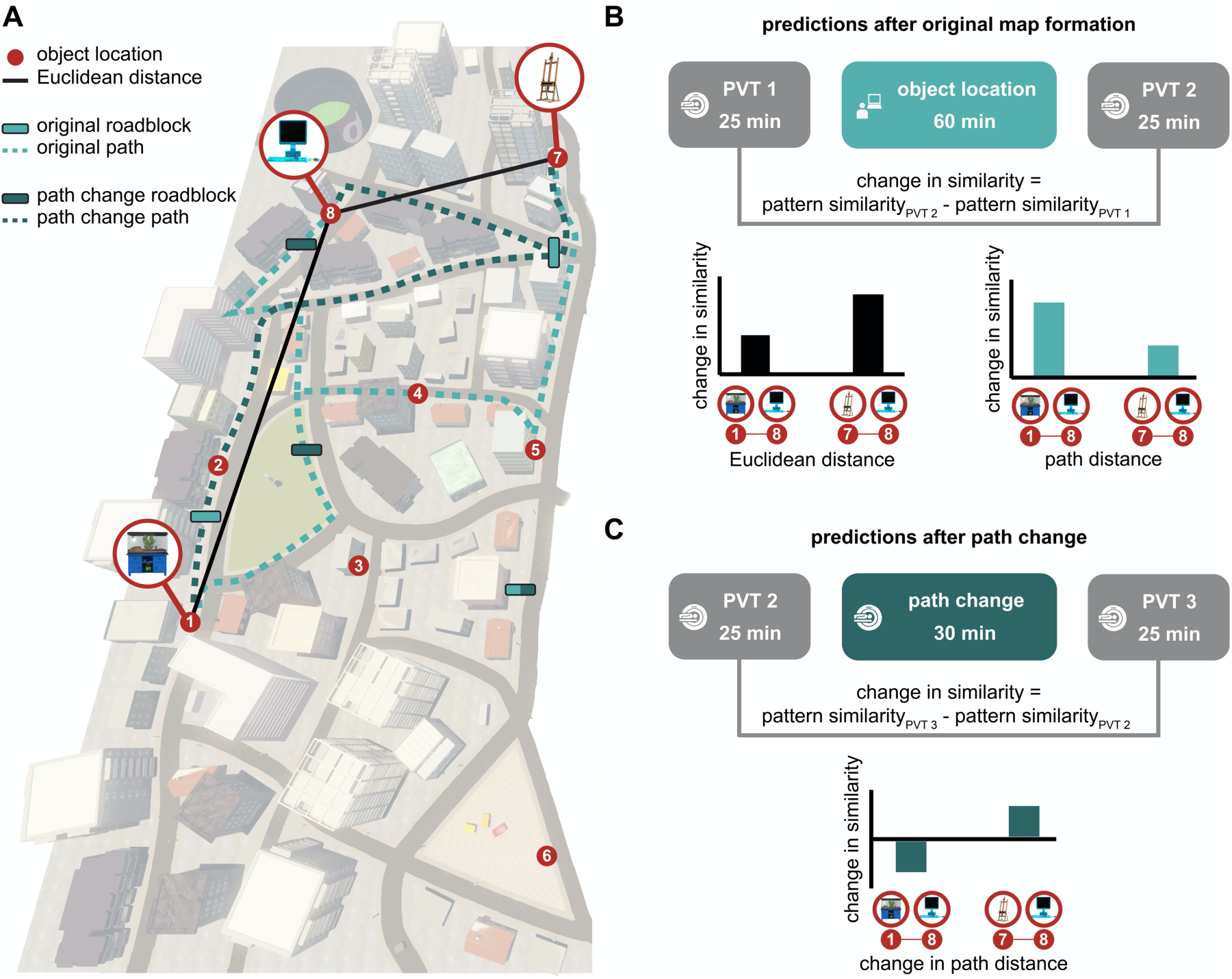
Disentangling Euclidean and path distances. A. Birdview of the virtual city, including numbered object-locations and three examples of objects. Roadblock locations during the object-location task are indicated by light bars, and roadblock locations during the path change task are indicated by dark bars. One roadblock was used in both tasks (light/dark bar). Two examples of object-to-object Euclidean distances are indicated by black lines. The dashed lines represent two examples of shortest object-to-object path distances during the object-location task (light) and path change task (dark). B. Predictions based on the Euclidean and path distances of the original map, during the object-location task. Examples are shown for three object-locations. If Euclidean distance is represented, we expect that objects 7 and 8 show a higher change in similarity when subtracting pattern similarity of PVT 1 from PVT 2, than objects 1 and 8, because the Euclidean distance is lower between 7 and 8 than between 1 and 8. However, because of the roadblocks, the path distance between 7 and 8 is higher than between 1 and 8. Therefore, if path distance is represented, we expect a lower similarity between 7 and 8 than between 1 and 8. C. Predictions based on the change in path distance during the path change task. Path distance between object 1 and 8 increases in the path change task, so we expect a decrease in pattern similarity when subtracting PVT 2 from PVT 3. We expect the opposite for object 7 and 8, where the path distance decreases in the path change task.

To assess neural representations of Euclidean and path distances, we measured neural pattern similarity between object-pairs, similar to Deuker and colleagues^13^. To this end, we analysed the fMRI data recorded during the PVTs using representational similarity analysis (RSA). RSA allowed us to calculate the neural similarity between object-pairs. We subtracted the neural similarity before the object-location task (PVT 1; Figure 1A) from the neural similarity after the first object location task (PVT 2; Figure 1A). This yielded the change in neural similarity between object-pairs that is induced by learning the object-locations in our navigation task. Since we expect that objects close in space are represented more similarly than objects far apart, we expect that this change in neural similarity scales with the distance between the object-pair (Figure 2B). To investigate differential representations of space in the left and right hippocampus, we performed ROI-analyses on both regions separately.

In order to investigate how changes in path distances affect neural representations, we furthermore calculated the change in neural similarity between object-pairs from before to after the second object location (path change) task (PVT2 to PVT 3; Figure 1A). We expected that for object-pairs with increased path distance the neural similarity would decrease, and vice versa for object-pairs with decreased path distance (Figure 2C). Therefore, we expected that there is a differential change of neural similarity for object-pairs with increased path distance compared to object-pairs with decreased path distance. Furthermore, we expected that the path distance change would affect hippocampal representations in persons who are sensitive to egocentric features (i.e., response learners) more that it would in persons who are sensitive to allocentric features (i.e., place learners).

### 2.3 Navigation tasks

#### 2.3.1 General set-up of the environment

Participants completed the three navigation tasks in the same virtual environment (Figure 1A). All active components of the environment and the navigation tasks were programmed and created in Unreal Development Kit 3. The environment had a length of ca. 19500 virtual distance points and a width of ca. 8760 virtual distance points. Throughout all tasks, participants were only able to navigate on the streets of the environment. Participants navigated with a constant speed of 600 virtual distance points per second (it took around 33 seconds to navigate from the South to the North end of the city along the most direct path). Participants completed all tasks from a first-person perspective and with a viewpoint-height of 90 virtual distance points. In order to move forward, participants could use the upward arrow-key on the keyboard and to rotate they could use the left and right arrow key. In the MRI scanner, participants indicated their responses with a button box with their right hand. Here, the middle button was used to move forward and the left and right button was used to rotate.

There were eight goal-locations in the virtual environment. All goal-locations were marked by a black box and were placed on the street grid of the virtual environment (Figure 1B, C). Each black box had the dimensions of 64−64−64 virtual distance points. The goal-locations remained constant across all three tasks. Euclidean distance between goal-locations varied between 2760 and 13242 virtual distance points. Path distance between goal-locations varied between 3362 and 17990 virtual distance points up until the path change task. For the path change task, path distance between goal-locations varied between 3117 and 20873 virtual distance points (see Figure 3 for an overview of all Euclidean and path distances between goal-locations). There were three roadblocks in the virtual environment. Two of them changed location between the object-location task and the path change task (Figure 2A). This resulted in changes in path distance ranging from −8517 to 7821 virtual distance points. Participants could not continue navigating on the street when encountering a roadblock and had to change direction to find another path. Roadblocks had an approximate height of the viewpoint of the player and a width of 56 virtual distance points.

**Figure 3.**
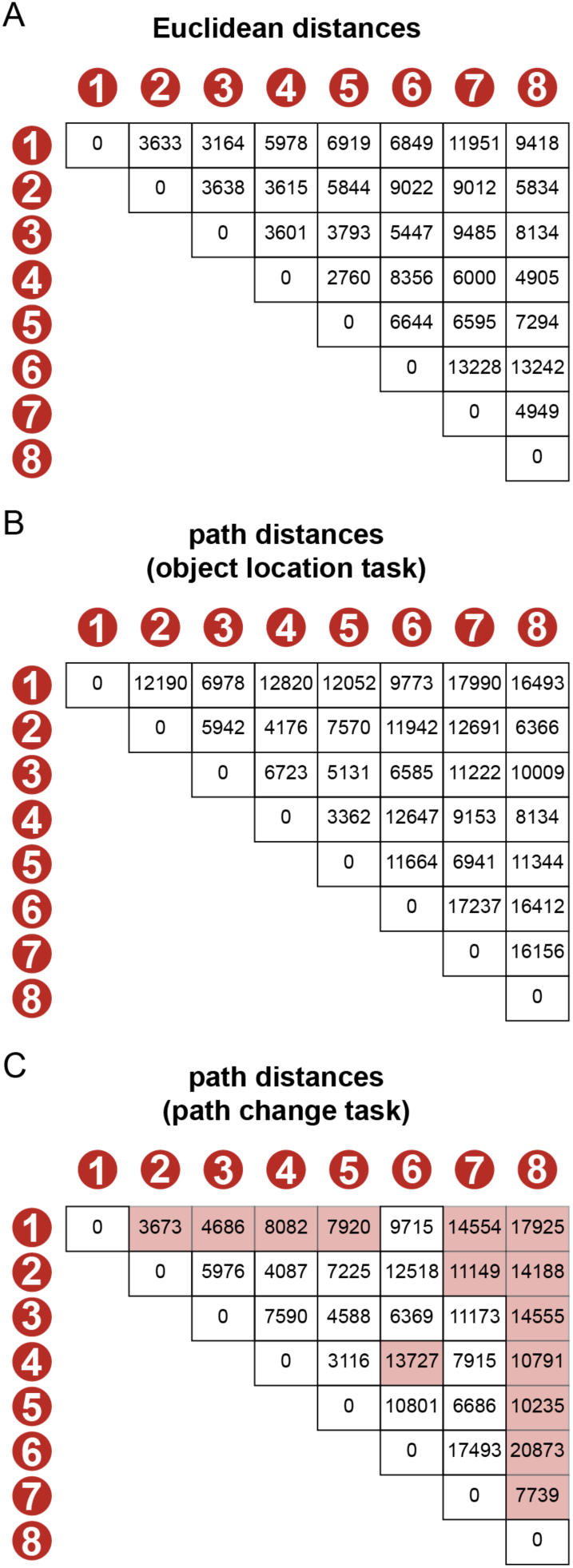
Distances between goal-locations, in virtual distance points. A. Euclidean distances between objects. These distances were the same during the object-location and path change task. B. Path distances between objects during the object-location task. C. Path distances during the path change task. Distances that were subject to a meaningful change (1000 virtual distance points) are marked pink.

#### 2.3.2 Training task

The aim of the training task was for participants to become familiar with the virtual environment. Particularly, we wanted participants to learn the eight goal-locations and the shortest route between them. For this purpose, trials in the training tasks consisted of participants navigating from one goal-location to another (Figure 1C). When reaching the correct goal-location, participants received feedback about whether they took the shortest possible route. There was a total of 112 trials. The order from which goal-location participants had to navigate to the next goal-location was pseudo-randomised. Every goal-location was the starting point and destination point an equal amount of times over the course of the experiment (in total 14 times). Furthermore, from every goal-location, participants navigated an equal amount of times to every other goal-location (in total two times). A navigation trial consisted of the following steps:

1. Participants saw instructions on the screen to go to the next box and find the shortest possible route for 1000ms. The target box turned multi-coloured for the duration of the trial.
2. Participants could start to navigate freely in order to find the target box. During navigation, participants received feedback about the Euclidean distance between their current location and the location of the target box. This feedback resembled a Wifi-signal. The closer the participant was to the target box, the stronger the signal became (i.e. the more bars appeared, with a maximum of eight bars). The signal was scaled to the maximum Euclidean distance of the experiment.
3. Once participants walked into the target box (reached the destination), they received feedback about whether or not they had taken the shortest possible route. This feedback was presented for 2000 ms.
4. The trial ended and the target box would become the start location of the next trial.

The 112 navigation trials were split into four blocks (of 28 trials). After each block, we tested participants’ knowledge about the locations of the eight boxes. All blocks were removed from the environment for this recall phase. Participants were asked to remember the location of every box. Participants could indicate their answers by pressing a button to signal that they were at a box-location. If they remembered the location of a box to be within a radius of 500 virtual distance points (about 2.6% of total map length South-to-North), participants received positive feedback. Otherwise, the box would appear at its correct location in multi-colour. Before continuing with the task, participants had to walk to the miss-located box and touch it. During the recall phase, participants saw the number of locations they still had to find.

Out of the 32 participants, 28 completed all four blocks of the task. As the task was self-paced, we stopped the task for four participants because the allotted lab time ran out. Two of these participants completed two blocks and the other two completed three blocks. There were no timing issues with the navigation tasks on the following day. On average the duration of the training task was 93 minutes (SD = 14 minutes). Participants got faster during the task, with the first block taking on average 26 minutes (SD = 8 minutes) and the last block taking on average 20 minutes (SD = 3 minutes). From block to block, participants (n = 28) also increased the number of trials in which they took the shortest possible route (first block: mean = 34%, SD = 14%, second block: mean = 47%, SD = 12%, third block: mean = 60%, SD = 12%, fourth block: mean = 61%, SD = 16%; Figure 4A). In the first block, participants took on average the shortest route in 34% (SD = 14%) of trials. In the last block, participants took on average the shortest route in 61% (SD = 16%) of trials. Over the whole task, participants took on average the shortest route in 49% (SD = 12%) of trials. Lastly, participants (n = 28) improved their knowledge about box-locations from block to block, reflected in an increase in correctly placed boxes (first block: mean = 53%, SD = 17%, second block: mean = 70%, SD = 20%, third block: mean = 88%, SD = 14%, fourth block: mean = 89%, SD = 17%; Figure 4B).

**Figure 4.**
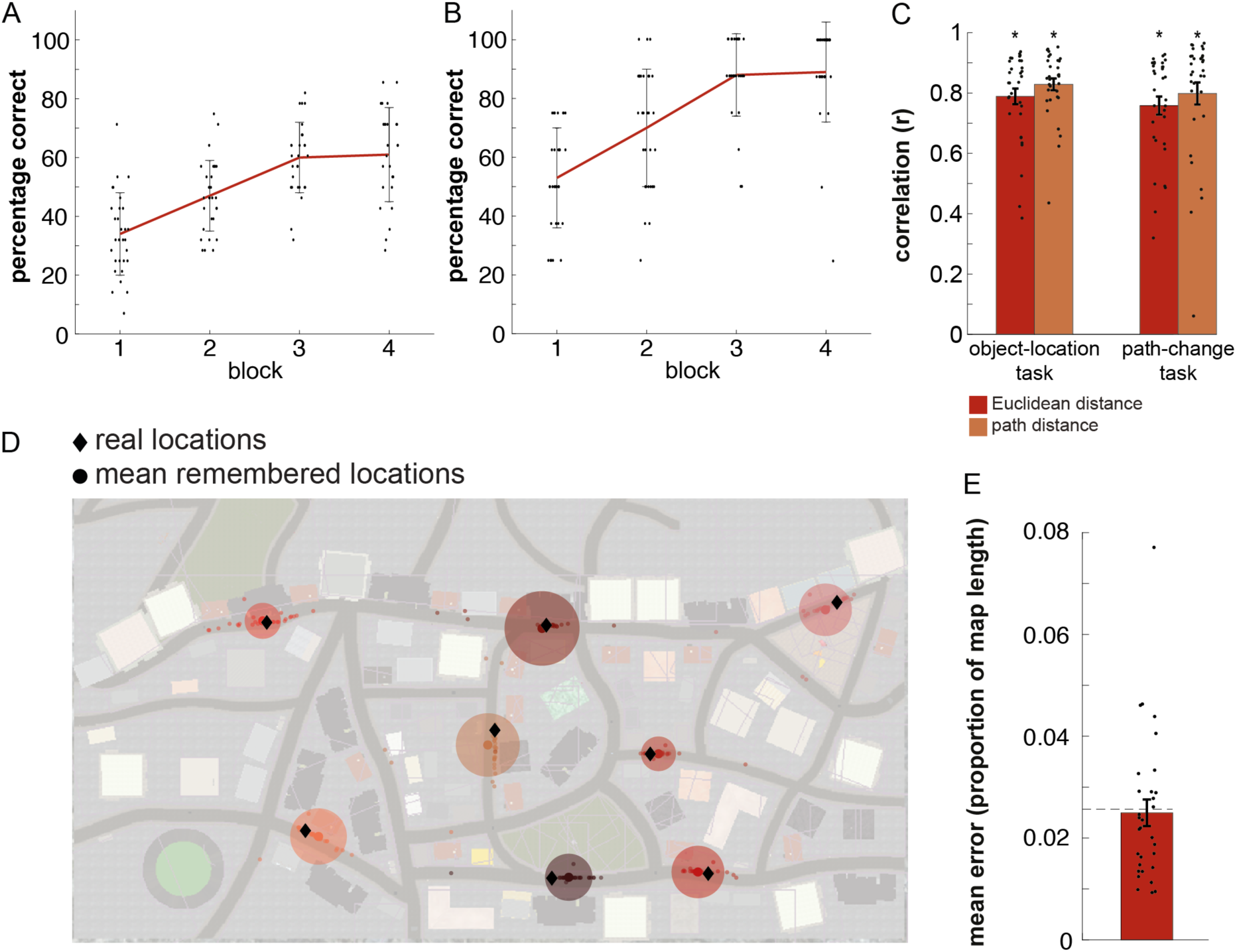
Behavioural results. A. Mean percentage of trials in which participants took the shortest route during the four blocks of the training task. Errorbars denote standard deviations. Small dots are individual data points. B. Mean percentage of correctly placed boxes at the end of the four blocks of the training task. Errorbars denote standard deviations. Small dots are individual data points. C. Results of the distance recall tasks before (left) and after (right) the path-change task. Errorbars denote SEM, dots are individual data points. D. Results of the bird-view placement task. Diamonds indicate the true locations, and large dots the mean placed locations. Circles indicate standard deviations around the mean placed location. Small dots are individual data points. E. Mean error of the bird-view task, as a proportion of total map length. The dashed line indicates the error radius of 2.6% that was allowed in the training task. Errorbars indicate SEM, dots are individual data points.

#### 2.3.3 Object-Location task

The object-location task was similar to the training task. Again, participants had to navigate from goal-location to goal-location and find the shortest possible route between them. A key difference however, was that every location/box was associated with one object, respectively.

Participants were presented with the object for 2000 ms, once they arrived at the target box. In total there were 112 trials – the same number of trials as for the training task. The rules for the pseudo-randomisation were the same as for the training task. There were no recall blocks for box locations. The task was split into two halves. In the first half, participants received feedback about their distance to the target-box while navigating (identical to the training task). In the second half, participants saw the object of the target box as a cue instead of the distance feedback (Figure 1C). Participants were instructed to go to the location of that object. With this manipulation, we aimed to encourage participants to learn the object-locations, instead of relying solely on the distance signal. For each participant, the eight objects were randomly selected from a pool of twelve objects (Figure 1E). Object-location associations were randomised across participants. All participants (n = 32) completed the object-location task. On average the task took 64 minutes (SD = 10 minutes). The first half of the task took on average 35 minutes (SD = 5 minutes) and the second half 29 minutes (SD = 6 minutes). Participants improved over the task in finding the shortest possible route (first half: 62% of trials (SD = 13%), second half: 87% of trials (SD = 14%).

#### 2.3.4 Path change task

The path change task was similar to the second half of the object-location task. In fact, the structure of the tasks and trials was identical to the second half of the object-location task (Figure 1C). The key difference was that the location of two of the three roadblocks had changed (Figure 2A). Therefore, the shortest route could change between the objects/boxes. In order to receive positive feedback, participants had to find these new routes. In total, there were 56 pseudo-randomised trials. Rules for the pseudo-randomisation were the same as for the training and object-location task. This resulted in seven trials per box/object as starting point and destination. Participants navigated from every object/box to every other object/box once during the path change task. All participants (n = 1. 32) completed the path change task in the MRI scanner. On average the task took 30 min and 23 sec (SD = 2 min and 55 sec). Participants took the shortest route on 74% (SD = 9%) of trials.

### 2.4 Testing knowledge about the map

We tested participants’ knowledge about the map on three different occasions. We asked participants to estimate the Euclidean and path distance for every object-pair after the object-location task and after the path change task, respectively (distance recall task). Lastly, we asked participants at the end of the whole experiment to indicate the location of every object on a map of the city (bird-view placement task).

For the distance recall task, we first asked participants to estimate all Euclidean distances between object-pairs and then to estimate all path distances between object pairs (in total 28 trials per distance estimation). The order of object-pairs was completely randomised across participants. For both estimations, we instructed participants to indicate the distance by entering a number between 0 and 100. We told participants that 0 would mean both objects share the same location, whereas 100 represents the longest possible distance between two objects. The task was completely self-paced. The task was programmed in neurobs Presentation (version 16.4, www.neurobs.com/presentation). We scored participants’ performance by correlating the estimated distances with the actual distances between object-pairs.

For the bird-view placement task, participants were presented with the map of the street-layout of the city. All buildings and landmarks had been removed from the picture. The street-layout was overlaid with a regular grid to help participants to indicate the locations. Next to the picture of the street-layout, participants saw the eight objects with a number next to every object. We asked participants to indicate the location of each object by writing its corresponding number onto the map (paper-pencil format). We scored participants’ performances by calculating the mean placement error (distance) between the recalled location and the actual location of every object.

### 2.5 Picture viewing tasks

In order to measure changes in neural pattern similarity, participants completed three (identical) picture viewing tasks (PVTs) in the MRI scanner. During a picture viewing task, pictures of the objects that participants encountered in the virtual city appeared on the screen one at a time. In order to keep participants’ attention high during the PVTs, participants had to perform an oddball task. The oddball was the picture of an object (a bathtub) that did not appear in the rest of the experiment. Participants had to press one of two buttons if the oddball object was presented (catch trials) and the other button if any other object was presented (regular trials). Button-contingencies were randomised across participants, but were always done with the right index and middle finger. This oddball task was orthogonal to later analyses of the PVTs. Participants (n = 32) performed at ceiling on the cover task in all PVTs (PVT 1: mean = 96.1%, SD = 17.5, PVT 2: mean = 98.7%, SD = 1.8, PVT 3: mean = 98.1, SD = 4.4).

The order of the objects was pseudo-randomised for every participant, but held constant across picture viewing tasks of the same participant. All eight objects that a participant encountered in the object-location and path change task were presented 20 times, respectively during a PVT. An object presentation took 2000 ms and the object was presented at the centre of a white screen. The order of objects was pseudo-randomised and each object was presented once within mini-blocks of eight regular trials. We made sure that the same object was never presented twice in a row. We added 32 catch-trials in which the oddball was presented. Hence, in total each PVT consisted of 192 trials. The oddball probability was constant across the experiment (oddball followed every regular object 4 times, probability was therefore 20%). Similarly, catch trials were equally distributed over the PVT with the same probability of appearing within every 20% of the task.

After every trial, a black fixation cross appeared at the center of the screen. ITIs could be either 2.5s, 4s or5.5s. The order of the ITIs was pseudo-randomised in a manner that all ITIs appeared equally often across all trials (the ITI of a catch trial was randomly chosen). Furthermore, the average ITI within every 20% of the task was not allowed to deviate more than one standard deviation of the average ITI of the whole PVT. The purpose of these pseudo-randomisations was to ensure that there was no temporal bias of ITI duration between objects or within the PVT. After every block of 60 non-catch trials, participants had a 20 second break, in which they received feedback about their performance on the previous block and a reminder of the button-contingencies. All PVTs were programmed in and presented with neurobs Presentation (version 16.4, www.neurobs.com/presentation).

### 2.6 Navigation ability assessment tasks

#### 2.6.1 Santa-Barbara Sense of Direction Scale

The Santa Barbara Sense of Direction Scale (SBSOD) is a self-referential questionnaire about general navigational abilities and sense of direction, with 15 Likert-type questions^22^. Participants completed the questionnaire on a computer, and could take as much time as they wanted. Under each question was a 7-point scale reaching from ‘strongly agree’ (1) to ‘strongly disagree’ (7). The final score was between 1 and 7, with a higher score reflected higher self-reported navigational abilities. On average participants had a score of 4.49 (SD = 1.18). The task was programmed in and presented with neurobs Presentation (version 16.4 www.neurobs.com/presentation).

#### 2.6.2 T-Maze task

The T-Maze task was adapted from Astur and colleagues^16^. The goal of the task was to identify tendencies towards ‘place learning’ and ‘response learning’. The task took place in a virtual room and was developed and programmed with Unreal Development Kit 3 (Unreal Engine 3, Epic Games, Inc.). We placed a T-shaped platform in the room (Figure 1D). Around the platform, we placed several landmarks in the room in order for participants to orientate themselves. We placed a multi-coloured box in either arm of the T-Maze, respectively. We instructed participants that one of the two boxes was rewarded, whereas the other was not. At the beginning of a trial, participants were placed at the bottom of the T-Maze. Participants could freely navigate on the platform with the help of the arrow keys (akin to the navigation tasks) and additionally rotate with the mouse. Once participants arrived at one of the two boxes, they received either positive or negative feedback for 2000 ms. If participants walked to the rewarded box, they were presented with a green happy smiley, otherwise they were presented with a red sad smiley. After the feedback, participants were presented with a black fixation cross on a white screen for 5000 ms and the next trial started.

In total, there were 10 trials. Two out of the 10 trials were probe trials. During these probe trials, the T-platform was rotated 180°. The idea here was that participants could either walk to the same location in reference to the room (indicative of allocentric place learning) as in previous rewarded trials or make the same turn as in previous rewarded trials (indicative of egocentric response learning). As we did not want to reward one strategy over the other, both choices were rewarded in probe trials. The first probe trial was either after 3, 5, or 7 normal trials (randomly chosen across participants), the second probe trial was always the last trial of the task^16^. At the beginning of the task, participants were placed at the bottom of the T-Maze and were instructed to look around and get familiar with the environment for 20 seconds. In that time, participants could not move away from the position they were in, but were able to rotate.

Out of the 32 participants 13 were classified as place learners, 17 were classified as response learners and 2 were classified as mix learners (based on inconsistent choices across the two probe trials). We tested for differences between place and response learners for performance on the behavioural tasks, as well as for neural representation of distances. Mixed learners were excluded from these comparisons.

### 2.7 Behavioural analyses

Behavioural signatures of a cognitive map of distances were assessed using distance recall tasks. We correlated estimated distances with actual distances between object-pairs. We used one-sample t-tests to test for an effect of the scores of all distance recall tasks after applying Fisher transformation to the correlation coefficients. As control analyses we correlated the estimated Euclidean distance with the real path distance and vice versa, the estimated path distance with the real Euclidean distance after the object-location task and path change task, respectively. We used a paired-sample t-test to test whether memory scores were significantly higher than their control counterparts. We analysed performance on the bird-view task by calculating the mean placement error (Euclidean distance) between the recalled location and the actual location of every object. We used a one-sample t-test to test for an effect of the scores on this task.

We further correlated both mean score of the distance recall task (after Fisher transformation) and the bird-view placement task with self-reported navigational abilities (SBSOD-scores) across participants. To test for behavioural differences between place and response learners in representing distances, we used two-sample t-tests on all scores of the distance recall task (after Fisher transformation) and the bird-view placement task. We also used a two-sample t-test to test for differences regarding self-reported navigational abilities (SBSOD-scores).

### 2.8 fMRI Analysis

#### 2.8.1 MRI Image acquisition

Functional T2*-weighted and anatomical images were acquired on a Magnetom Prisma or PrismaFit 3 Tesla magnetic resonance tomograph (Siemens, Erlangen, Germany) with a 32-channel head coil. Due to the extensive nature of the study and booking load of the PrismaFit scanner, we decided to collect four out of the 32 data-sets on the Prisma scanner. Both sessions within each participant were scanned on the same scanner. Functional images were acquired with a 4D multiband sequence with 84 slices (multi-slice mode, interleaved), TR = 1500 ms, TE = 28 ms, flip angle = 65 deg, acceleration factor PE = 2, FOV = 210 x 210 x 168 mm and an isotropic voxel size of 2 mm. An anatomical image of the brain was acquired, using a T1 sequence (MPRAGE) with TR = 2300 ms, TE = 3.03 ms, flip angle = 8 deg, FOV = 256 x 256 x 192 mm and an isotropic voxel size of 1 mm. For 29 participants on day 1 and 28 on day 2, at the end of each scanning session two separate phase and magnitude images were acquired in order to correct for distortions with a gradient field map (multiband sequence with TR = 1020 ms, TE = 10 ms, flip angle = 45 deg and a voxel size of 3.5 x 3.5 x 2.0 mm).

#### 2.8.2 fMRI preprocessing

Functional images of the three functional runs (one per PVT) were preprocessed with help of the FSL toolbox (version 5.0.4, http://fsl.fmrib.ox.ac.uk/fsl/fslwiki/). Motion correction (three rotation and three translation estimations) and a high pass filter (cut-off: 100s) were applied to the images. The anatomical scan of each participant was downsampled to the voxel size of the functional scans (2 mm isotropic). In order to have a common reference space for the first-level analysis, all functional scans were linearly registered to the down-sampled anatomical scan.

After preprocessing, we excluded participants from further analysis of a functional run based on the following criteria: 1. No appropriate responses during the PVT (minimum inclusion criteria was pressing both possible buttons, no participant was excluded). 2. More than 10% of the volumes of a functional run had movement above 3 mm (two participants excluded from analyses of PVT 2 and an additional participant excluded from PVT 3). Taken together, 30 participants (12 place learners, 16 response learners and 2 mixed) were included for analysing changes from PVT 1 to PVT 2 and 29 participants (12 place learners, 15 response learners and 2 mixed) were included for analysing changes from PVT 2 to PVT 3.

#### 2.8.3 First-level fMRI analysis

We used representational similarity analysis (RSA) to measure changes in neural similarity between object-pairs as a proxy for a cognitive map (Kriegeskorte et al., 2008; Deuker et al., 2016). More specifically, we calculated neural pattern similarity between object-pairs by correlating the neural activation pattern of every object with every other object, both before (PVT 1) and after (PVT 2) the object-location task (Figure 2B), and before (PVT 2) and after (PVT 3) the path change task (Figure 2C). Subsequently, we subtracted the pattern similarity of PVT 1 from PVT 2, and of PVT 2 from PVT 3, to test whether change in pattern similarity scaled as a function of Euclidean distance and/or path distance. Furthermore, we added a term for integration between Euclidean and path distance. We did this because Euclidean and path distance are never completely decoupled (r = 0.57), and path distance is highly related to the average travel duration between objects in our design (object-location task: r = 0.99, path change task r = 0.96). See the section on ROI analyses for more details.

##### General Linear Model

We estimated object-specific activation by modeling the onset and duration for each object in a GLM (one GLM per PVT). To account for the other events during the PVTs, we set up additional regressors. All catch trials (oddball trials) were modeled in a single separate regressor. Additionally, we modeled button presses with the index and middle finger in two separate regressors with a stick function. The beginning of the task was modeled with the duration from start time of scanning until the first object presentation. The end of the task was modeled with the duration from the end of the last object presentation until end of scanning. Furthermore, all block breaks were modeled in one regressor with the duration from the end of the last object presentation before the break until the first object presentation after the break. Lastly, we accounted for movement by adding six movement parameters (estimated during preprocessing) and added an additional regressor for each volume that exceeded a movement of 3 mm (on average 1.7 volumes per run SD = 7.6).

##### ROI-Analysis

We performed analyses on anatomical masks of the left and right hippocampus. Both hippocampus masks were based on the Harvard-Oxford subcortical structural atlas (https://fsl.fmrib.ox.ac.uk/fsl/fslwiki/Atlases). Included voxels had to fulfill two criteria: a minimum probability of 25% to be hippocampus voxels and no higher probability to belong to a region other than the hippocampus. We only included grey-matter voxels into our masks. As these masks were in MNI space, we translated the object specific parameter estimates of every PVT into MNI space. To estimate neural similarity between object-pairs, we correlated the parameter estimates of every object with the parameter estimates of every other object across all voxels within the ROI mask (Spearman’s correlation coefficient). Subsequently, we could then subtract these correlation values of a pre-navigation-task PVT from a post-navigation-task PVT (e.g. 2-1, 3-2) as index for change in neural similarity. We were interested how these changes in neural similarity were affected by distances in the map. In other words, we wanted to test how well the resulting object-to-object correlations fit with our distance predictions (Euclidean, path, and integration). We performed multiple linear regression on the object-to-object correlations within each ROI, with the different distances as model predictors. Multiple linear regression allowed us to account for shared variance between the different distance measurements.

##### Distances as Model Predictors

We used distances between objects as model predictors for the ROI-analysis described above. To index offline map representation of distances in the hippocampus, we first measured the effect of Euclidean and path distance on change in neural similarity from PVT 1 to PVT 2. We performed a median split on object-pairs based on their Euclidean and their path distances, respectively. We set the weight for low distances to 1 and for high distances to 2. We entered as predictors the weight for Euclidean distance and the weight for path distance. We also tested the effect of the combined Euclidean and path distance by adding an integration term and a constant term. The integration term was the product of the Euclidean prediction and the path prediction. This way, the highest weight was given to object-pairs that have a high Euclidean and a high path distance and the lowest weight to object-pairs with a low Euclidean and low path distance. As a result, we obtained a beta estimate for three distance predictors per ROI per participant.

As a second step, in order to measure how changes in path distances affect hippocampal representations, we assessed how changes in similarity from PVT 2 to PVT 3 were affected by changes in path distances. We only included object-pairs that experienced a meaningful change in path distance. We defined a meaningful change as a distance change higher than 1000 virtual distance points (Figure 3C). We chose this cut-off as participants received positive feedback about placing the goal-locations within a radius of 500 virtual distance points during the training (2 object-locations * allowed error of 500 virtual distance points). As a result, 15 object-pairs were included into the analyses. We performed a median-split on these object-pairs into a (relative) decrease group and a (relative) increase group. We then calculated the mean change in neural similarity across all object-pairs of one group. Subsequently, we subtracted the mean change in neural similarity of decrease object-pairs from increase object-pairs. As a result, we obtained one value per ROI per participant.

#### 2.8.4 Whole-brain Searchlight Analysis

We performed searchlight analyses on the whole-brain level (for voxelwise-output). We included into the searchlight a grey-matter mask, which was based on the participant-specific downsampled anatomical scan. The searchlight had a radius of 3 voxels and was thresholded to include a minimum of 30 grey-matter voxels. RSA analysis within a searchlight was analogous to the analysis within an ROI. We correlated the parameter estimates of every object with the parameter estimates of every other object across all voxels within each searchlight. We then subtracted these correlation values of a pre-navigation-task PVT from a post-navigation-task PVT (e.g. 2-1, 3-2). Within each searchlight, we tested how well the resulting object-to-object correlations fit with our distance predictors (Euclidean, path, and integration) using multiple linear regression. For each predictor, the resulting parameter estimate was written back into the center voxel of the searchlight. Before second-level analyses we spatially normalised the relevant outputs from the searchlight analyses to MNI anatomical space and afterwards smoothed them using a 4mm full-width at half maximum Gaussian kernel.

#### 2.8.5 Statistical Analysis

For the ROI analyses, we had one entry per participant per test and used a one-sample t-test when testing across all participants. We furthermore tested for differences between place and response learners using a two-sample t-test. We also performed correlations with the ROI betas and the SBSOD-scores across all participants and (where applicable) across all participants within the same navigation-strategy group (place or response learners).

For the searchlight analyses we used a one-sample permutation test when testing across all participants and a two-sample permutation test when testing for differences between place and response learners. Output images from the first level analyses were entered as input, as well as a whole-brain mask which only included grey-matter voxels where all participants had an entry. 10000 random sign flips were performed to estimate the null distribution. We used threshold-free cluster enhancement and corrected for multiple comparison with family-wise error rate (p < 0.05).

## 3. Results

### 3.1 Behavioural signatures of a cognitive map

We set out to test how the hippocampus represents Euclidean and path distances and furthermore how these representations adapt to changes in the environment. To answer this question on a behavioural level, we let participants estimate the Euclidean and the path distances between object-pairs, respectively, on a scale from zero to 100. As expected, all average scores were significantly greater than zero (Figure 4C). More specifically, after the object-location task, average correlation between estimated and real path distance was r = 0.83 (T_(31)_ = 21.91, p < 0.001) and average correlation between estimated and real Euclidean distance was r = 0.79 (T_(31)_ = 18.28, p < 0.001). Actual path and Euclidean distances correlated with r = 0.57. There was no significant difference between estimated path and Euclidean distance scores (T_(31)_ = 1.74, p = 0.09). To control that participants really learned Euclidean and path distance separately, we also correlated remembered path distance with the real Euclidean distance, and vice versa, remembered Euclidean distance with the real path distance. Distance memory scores were significantly higher than their control counterparts (path distance vs. control: T_(31)_ = 5.77, p < 0.001, Euclidean distance vs. control: T_(31)_ = 6.53, p < 0.001). After the path change task, the average correlation between estimated and real path distance was r = 0.80 (T_(31)_ = 14.76, p < 0.001) and the average correlation between estimated and real Euclidean distance was r = 0.76 (T_(31)_ = 16.14, p < 0.001). The path distance estimations were significantly better than the Euclidean distance estimations after the path change task (T_(31)_ = 4.21, p < 0.001). The same control analyses were applied here as after the object-location task. Distance memory scores after the path change task were significantly higher than their control counterparts (path distance vs. control: T_(31)_= 6.42, p < 0.001, Euclidean distance vs. control: T_(31)_= 4.90, p < 0.001).

In addition to the distance recall tasks, we presented participants with a bird’s-eye view of the virtual city, where they had to indicate the location of every object on the map. On average, participants had a distance error between the indicated and the real location corresponding to 2.5% (SD = 1.4% of the total map length; Figure 4D and E). This distance error was below the error radius of 2.6% which was set as the maximum error in the training task. Additionally, across all participants, the mean error did not differ from this maximum error (p = 0.281, nonparametric sign test). Taken together, the behavioural results indicate that participants could accurately remember object locations. They could furthermore separately recreate the Euclidean as well as path distances between them.

Furthermore, we found a positive relationship between self-reported navigational abilities (measured here with the SBSOD) and mean distance memory score of r = 0.56 (p = 0.001). Additionally, we found a negative correlation between SBSOD-scores and mean placement error in the bird-view task of r = −0.33 that reached trend level (p = 0.072). We also explored whether differences in navigational strategies were related to behavioural performance. There were no significant differences between response and place learners for any of the distance memory scores (Euclidean distance: p = 0.29, path distance: p = 0.25, Euclidean distance after path change: p = 0.40, path distance after path change: p = 0.63) or mean placement error in the bird-view task (p = 0.33). Lastly, there was also no significant difference between response and place learners for self-reported navigational abilities (p = 0.33).

### 3.2 Hippocampal signatures of Euclidean and path distances

In a next step, we set out to investigate hippocampal representations of Euclidean and path distances. Using representational similarity analysis (RSA), we calculated change in neural pattern similarity between object-pairs, by analysing fMRI data during picture viewing tasks (PVTs). Participants performed at ceiling level during PVT 1 (correct responses: mean = 96.1%, SD = 17.5), PVT 2 (correct responses: mean = 98.7%, SD = 1.8) and PVT 3 (correct responses: mean = 98.1%, SD = 4.4). We predicted that this neural similarity would scale with distance, i.e. that object-pairs with a low distance would have a higher neural similarity than object-pairs with a high distance.

#### 3.2.1 ROI analysis results

Based on our a-priori hypothesis, we used the left and right hippocampus as ROIs in the analyses. We used the three distance predictors, Euclidean distance, path distance, and the integration distance (see below), to test whether the neural similarity of object pairs in the left and right hippocampus scales with these distance types. It could be that the hippocampus does not distinctly represent Euclidean and path distances, but that it represents an integration of the two. It could thereby differentiate the extremes, and separate object pairs that have a high Euclidean as well as path distance from object pairs that have a low Euclidean as well as path distance. Since we systematically manipulated some paths using roadblocks, some object pairs have relatively high Euclidean but relatively low path distance, or vice versa. Therefore, we furthermore compared object-pairs that have a high Euclidean and high path distance with object-pairs that have a low Euclidean and low path distance to test whether the two distance types are integrated. Hence, the ‘integration distance’ was the product of the Euclidean and path distance.

##### Left hippocampus

We found an effect for all three predictors in the left hippocampus (Euclidean distance categories: T_(29)_ = 2.3768, p = 0.048, path distance categories: T_(29)_ = 2.0255, p = 0.052 approaching significance, integration of Euclidean and path distance categories: T_(29)_ = −2.5570, p = 0.048, corrected for multiple comparisons using a Bonferroni-Holm correction (Figure 5A). The direction of the integrative effect of Euclidean and path distances, as measured by the integration term was as expected. The lower the integrated Euclidean and path distance was between object-pairs, the higher the increase in similarity. However, the directions of the separate effects for the Euclidean and path distance were unexpected, with a higher increase in similarity for high distances than low distances.

**Figure 5.**
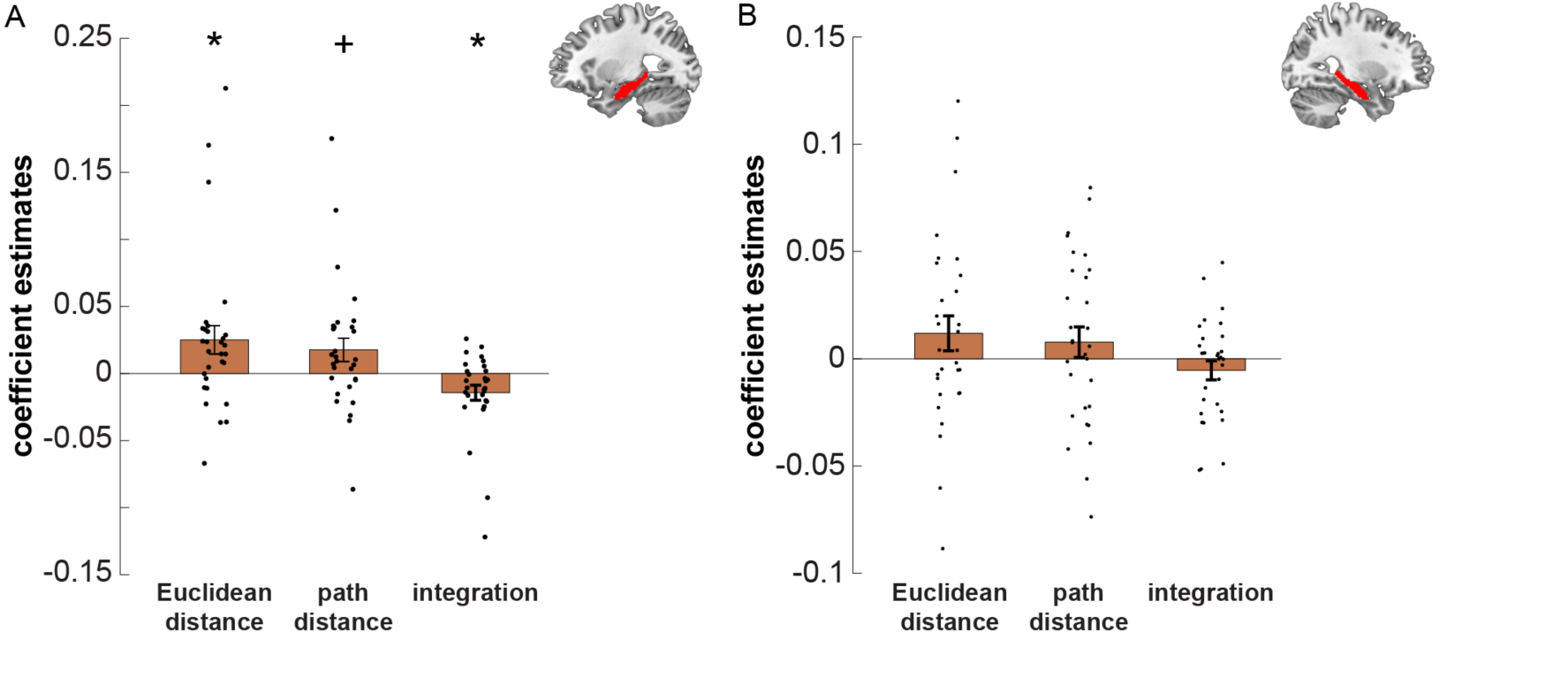
ROI analysis results for left and right hippocampus. Results of the ROI analyses for representation of Euclidean and path distances and their integration after the object-location task, using representational similarity analysis. Coefficient estimates are the result of the multiple linear regression analysis we performed on the object-to-object correlations within each ROI, with the Euclidean distance, path distance, and their integration as predictors. The change in neural similarity is shown from before (PVT 1) to after (PVT 2) the object-location task. Bars indicate mean ± SEM. A. Results in the left hippocampus. We found a significant effect for Euclidean distance and integration, and a trend effect for path distance, using multiple linear regression. B. Results in the right hippocampus. We found no significant effects. ^+^p < 0.1 (trend), * p < 0.05.

Furthermore, Euclidean distance representations in the left hippocampus, but neither path distance representations nor the representation of the integration of Euclidean and path distances, correlated with SBSOD-scores — an indicator of self-reported navigational abilities (Euclidean distances: r = −0.385, p < 0.05; path distances: r = −0.310, p = 0.096; integration Euclidean and path distances: r = 0.318, p = 0.087).

Based on the allocentric nature of Euclidean distances and the egocentric nature of path distances, we additionally explored differences in hippocampal representations between place learners (egocentric navigators) and response learners (allocentric navigators; both groups were classified by an independent T-maze test). We found no differences between place and response learners for neural representations that scale with distance in the left hippocampus (Euclidean distance categories: p = 0.613, path distance categories: p = 0.752 integration of Euclidean and path distance categories: p = 0.659).

##### Right hippocampus

There was no effect for Euclidean or path distances in the right hippocampus, when testing across all participants (Euclidean distance category: p = 0.156, path distance category: p = 0.287, integration of Euclidean and path distance category: p = 0.228; Figure 5B). However, we observed a trend effect for differences between place and response learners for effects of Euclidean distance categories (T_(26)_ = −1.9786, p = 0.059) and the integration of Euclidean and path distance categories (T_(26)_ = 1.9016, p = 0.068), but not path distance categories (p = 0.109, Figure 6A). Post-hoc tests reveal that these differences were probably driven by the response learners. Response learners showed significant or trend effects (Euclidean distance categories: T_(15)_ = 2.2957, p = 0.037, path distance categories: T_(15)_ = 1.9668, p = 0.068 integration of Euclidean and path distance categories: T_(15)_ = −2.1259, p = 0.051, Figure 6A), whereas the place learners did not (Euclidean distance categories: p = 0.65, path distance categories: p = 0.67 integration of Euclidean and path distance categories: p = 0.59, Figure 6A).

**Figure 6.**
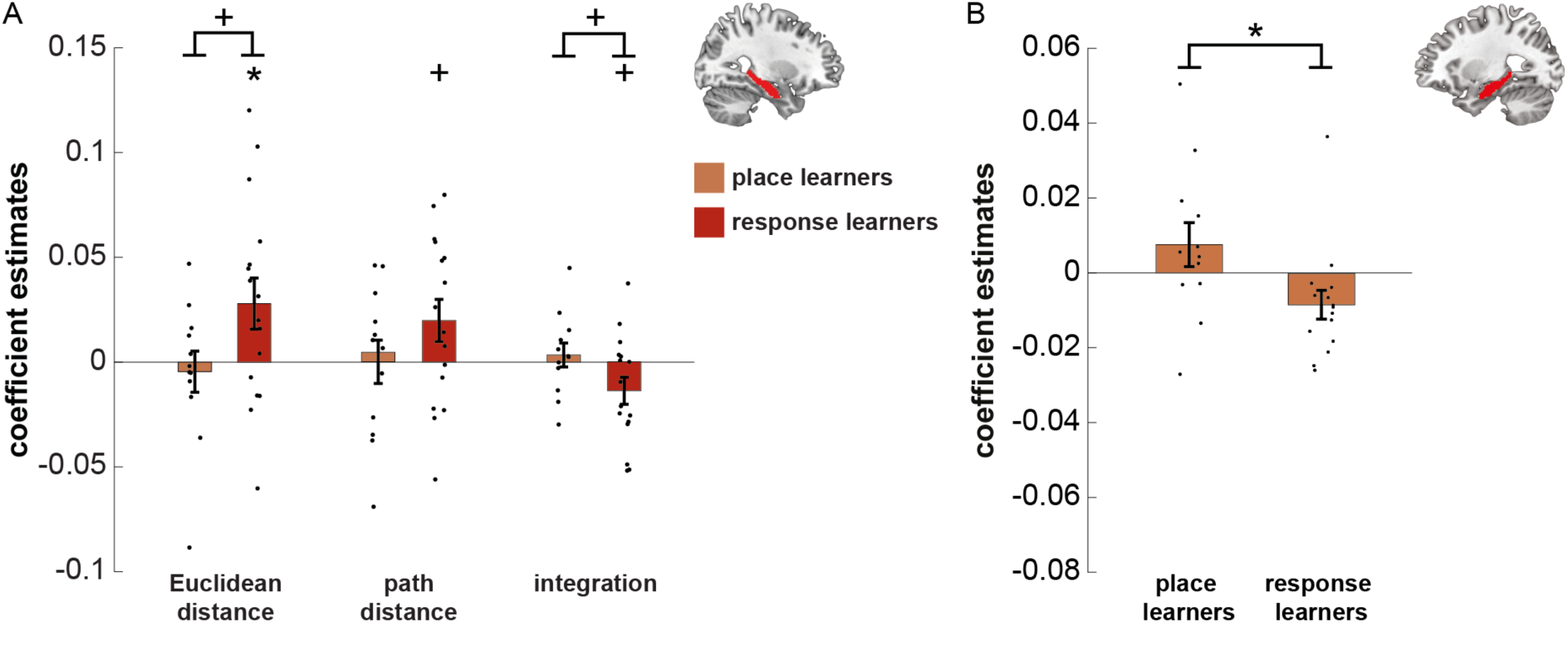
Post-hoc results of differences between place and response learners. A. Results of the post-hoc analyses on the right hippocampus after the object-location task. Coefficient estimates are the result of the multiple linear regression analysis we performed on the object-to-object correlations within each ROI, with the Euclidean distance, path distance, and their integration as predictors. We found trend differences between the place and response learners in the right hippocampus, for Euclidean distance and integration. This was probably driven by the (trend) significant representations in response learners but not place learners. The change in neural similarity is shown from before (PVT 1) to after (PVT 2) the object-location task. Bars indicate mean ± SEM. B. Results of the post-hoc analyses on the left hippocampus after the path change task. We first estimated the change in neural similarity from pre (PVT 2) to post (PVT 3) of object pairs which path length increased versus which path length decreased. We then subtracted the mean change in neural similarity of decrease object-pairs from increase object-pairs. We found a significant difference between the place and response learners in representation of change in path distances. The change in neural similarity is shown from before (PVT 2) to after (PVT 3) the path change task. Bars show mean ± SEM. * p < 0.05

As only response learners showed any indication for a representation that scales with Euclidean distance in the right hippocampus, we explored whether those correlated with self-reported navigational abilities. Here, we found no significant correlations between distance representations in the right hippocampus and SBSOD-scores (Euclidean distance categories: p = 0.32, path distance categories: p = 0.61, integration of Euclidean and path distance categories: p = 0.37).

#### 3.2.2 Exploratory whole brain searchlight analysis results

We explored representations of path and Euclidean distances outside of the hippocampus. Searchlight analyses on grey-matter voxels at the whole-brain level revealed no significant representations of Euclidean distance or path distance that survived correction. We also found no differences between place and response learners for any distance category effects that survived whole-brain correction. Nevertheless, we show the non-thresholded effects in Figure 7.

**Figure 7.**
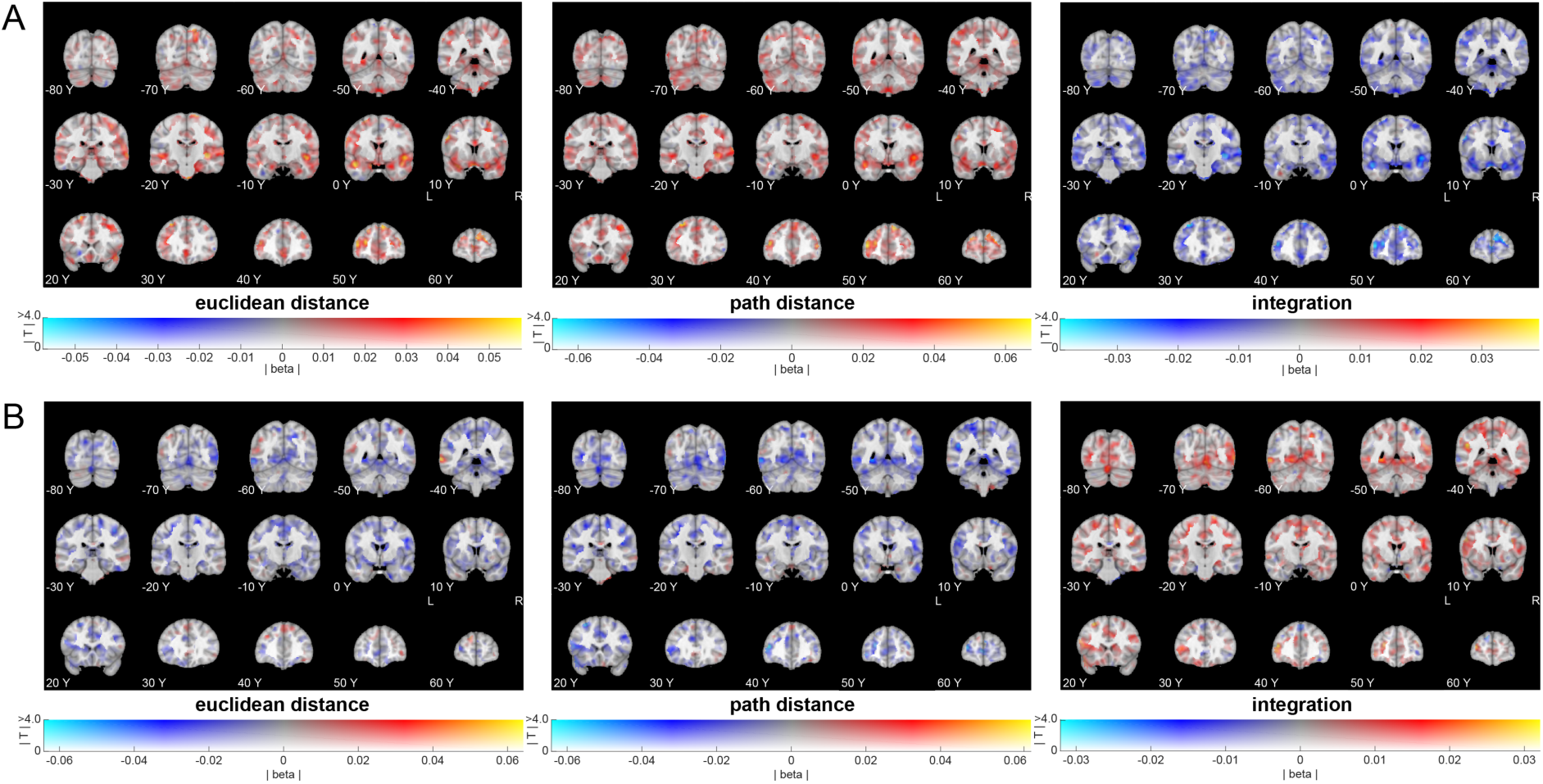
Whole brain searchlight results for effects from PVT1 to PVT2 (uncorrected whole brain effects). A. Whole brain effects of distance categories on change in neural similarity from before (PVT 1) to after (PVT 2) the object-location task. Left: effects of Euclidean distance categories. Middle: effects of path distance categories. Right: effects of integration distance categories. No effects survived whole-brain correction. B. Whole brain effects for differences between response and place learners in distance representations, from before (PVT 1) to after (PVT 2) the object-location task. Left: effects of Euclidean distance categories. Middle: effects of path distance categories. Right: effects of integration distance categories. No effects survived whole-brain correction. All images were created using a dual-coded design^24, 25^. This allowed showing both the mean beta coefficient (blue-red) and the T-stats (opacity). Y-coordinates are in MNI space.

Taken together, these results suggest that the left hippocampus forms representations that scale with integrated Euclidean and path distances. Furthermore, the Euclidean distance representations were related to self-reported navigational abilities. We found a trend effect that response learners and not place learners use the right hippocampus to represent Euclidean distances and an integration of Euclidean and path distances. Whole-brain analyses revealed no evidence for distance map representations outside of the hippocampus.

### 3.3 The effect of path distance changes on its hippocampal representations

The second main question of this study was whether a hippocampal map can react flexibly to changes in the environment. For this purpose, participants performed a path change task after PVT 2. Here, path distances between object-pairs had been altered compared to the object-location task, but Euclidean distances remained stable. (Figure 1A, 2C).

#### 3.3.1 ROI analysis results

Effects of path changes were assessed in the left and right hippocampus ROIs. To this end, object-pairs that were subject to a meaningful change in path distance (2 times the allowed error of 500 virtual distance points during the training) were median split into an ‘increased path distance’ and a ‘decreased path distance group’ (see Figure 2C and Methods for more details, and Figure 3C for actual distances). When testing across all participants, there was no effect of change in path length for neither the left (p = 0.462), nor the right hippocampus (p = 0.823). However, we found a significant difference in path change representation between place and response learners in the left hippocampus (T_(25)_ = 2.339, p = 0.028), but not the right hippocampus (p = 0.188). Post-hoc tests revealed that this effect was driven by response learners: they showed a significant effect of change in path length in the left hippocampus (T_(14)_ = −2.171, p = 0.048, Figure 6B) while place learners did not (p = 0.227, Figure 6B). The direction of the effect for response learners was as expected, with object-pairs in the path length decrease group showing a higher increase in neural pattern similarity than object-pairs in the path length increase group. We found no significant correlation between SBSOD-scores and the effect of change in path length across response learners (p = 0.839).

#### 3.3.2 Exploratory whole brain searchlight analysis results

We explored effects of path distance changes on representations outside of the hippocampus. Searchlight analyses on grey-matter voxels at the whole-brain level revealed no representations of changes in path length – neither across all participants, nor for differences between place and response learners. Nevertheless, we show the non-thresholded effects in Figure 8.

**Figure 8.**
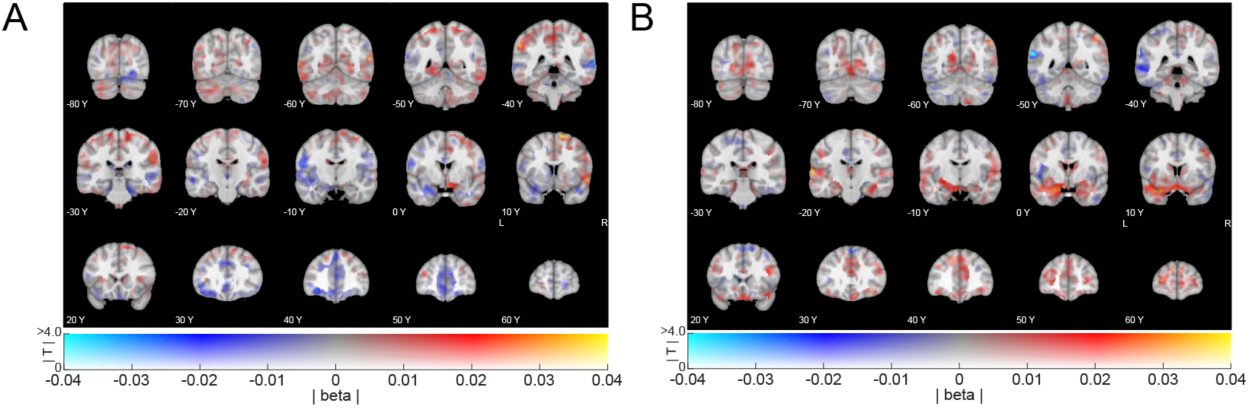
Whole brain searchlight results for effects from PVT2 to PVT3 (uncorrected whole brain effects). A. Whole brain effects of changes in neural pattern similarity from before (PVT 2) to after (PVT 3) the path change task, as a function of change in path distance. No effects survived whole-brain correction. B. Differences between path and response learners for representing change in path distance (measured as in A). No effects survived whole-brain correction. All images were created using a dual-coded design^24, 25^. This allowed showing both the mean beta coefficient (blue-red) and the T-stats (opacity). Y-coordinates are in MNI space.

Taken together, we found that response and place learners exhibit differential sensitivity of their hippocampal representations that scale with distance to changes in the environment. Specifically, the left hippocampal pattern similarity was sensitive to changes in path lengths. Place learners, however, showed no effect. Whole-brain analyses revealed no evidence for effects of path change outside of the hippocampus.

## 4. Discussion

The results of the current study demonstrate that participants acquired map-knowledge about relevant locations and the Euclidean and path distances between these places when navigating in a virtual environment. By associating goal-locations with objects, we were able to measure how Euclidean and path distances affect the neural pattern similarity between object-pairs as a proxy for a cognitive map representation. We found that the left hippocampus represents and integrates path and Euclidean distances. Our second main interest was whether such a map adapts to changes in the environment. Therefore, we changed path distances between object-pairs and tested how these changes affected neural pattern similarity. There was no effect of changes in path distance on hippocampal representations when testing across all participants. However, there was a significant difference between place and response learners, whereby response learners showed a significant effect of path distance changes, whereas place learners did not. This is in line with our expectation that place learners mainly represent allocentric features (Euclidean distances), which remain stable during the path-change task, and that response learners mainly represent egocentric features (path distances), which changed.

### 4.1 An integrative map in the hippocampus

Our results indicate that the left hippocampus forms a map that integrates Euclidean and path distance information (Figure 5A). This is in line with previous fMRI studies testing map representations in the hippocampus^12, 13^. Morgan and colleagues^12^ found evidence for both Euclidean and remembered distance travel time (comparable to path distances in our study) representations in the left hippocampus. Importantly though, these two distance types highly correlated (r = 0.9). In comparison, Euclidean and path distances had a lower correlation in our study (r = 0.57), since we deliberately designed paths in such a way to disentangle both types of distances. Furthermore, multiple linear regression allowed us to test the effects of either distance while simultaneously taking into account shared variance. Hence, our results provide more clear evidence that both types of distance information are represented and integrated by the hippocampus. This interpretation dovetails with the results of a previous study testing map representations in the hippocampus^13^. Here, the authors also found significant effects of remembered Euclidean distance, and for the interaction between remembered Euclidean distance and remembered distance travel time in the left hippocampus. A key difference of our design compared to theirs, is that our participants freely navigated from object to object instead of along a fixed path, making our design a more naturalistic reflection of real-world navigation. Nevertheless, given the consistency between our results and these fMRI studies^12, 13^, one speculation could be that our path distance measure actually relates to travel time between objects instead of spatial distance per se. The navigation speed from object to object was constant during our navigation tasks, hence, travel time highly correlated with path distance. Theoretically our results might therefore also reflect an interaction between time and space, instead of an interaction between Euclidean and path distance. This idea is in line with theories about cognitive mapping in the hippocampus^7, 26–31^. Accordingly, the hippocampus represents and processes spatial and temporal information in similar ways. In a broader sense, these theoretical accounts suggest that the hippocampus has general processing mechanisms that are domain unspecific. We cannot directly test this idea in the current study as we tested for distance representations in the spatial domain. However, our results demonstrate that the left hippocampus forms a map that integrates different spatial relationships between relevant locations. This, at the very least, is coherent with the idea that the hippocampus is able to form cognitive maps that entail and integrate a variety of information.

Interestingly, we found that Euclidean distances were represented in the opposite direction than expected: higher distances resulted in a higher increase in pattern similarity than lower distances. One possible explanation involves pattern separation. It could be that the hippocampus disambiguates overlapping representations, and therefore represents two close locations less similarly to be able to distinguish them. This is in line with other findings, where learning decreased similarity of hippocampal representations of overlapping routes^32^ or of similar stimuli^33, 34^. Lower neural similarity might prevent interference between similary memories to support ‘mnemonic repulsion’^32, 33^, which might be reflected in our finding that low Euclidean distances were represented less similarly than high distances.

Another aspect is that this finding points towards a relationship between self-reported navigational abilities and Euclidean distance representation. The positive, unexpected pattern similarity scaling effect with distance was negatively correlated with self-reported navigational abilities. This indicates that this unexpected pattern of distance representations is more pronounced the worse people (self-reportedly) navigate. It could be that in poor navigators, the hippocampus shows a stronger pattern separation effect to account for the worse navigational ablities. Furthermore, posthoc analyses indicate that response learners showed representations in the right hippocampus, but place learners did not. We speculate that response learners might benefit from stronger hippocampal activity to efficiently navigate an environment compared to place learners. It is, however, important to note that the number of participants in our study was probably too small to interpret these effects of inter-individual differences with high confidence. In future work, it might be beneficial to pre-screen potential participants and to categorise them into response and place learners to be able to test the effects of navigational abilities in a more direct manner.

### 4.2 How changes in the environment affect the hippocampal cognitive map of response learners, but not place learners

Being flexible and adaptive are key characteristics that are attributed to cognitive maps^1, 30, 31^. Nevertheless, human neuroimaging studies about the effects of changes in spatial environments on hippocampal representations are rare. To our knowledge, these studies are limited to studying these effects during active navigation^3, 10, 35–37^ (for an overview see ^6^). We wanted to expand on this by testing how changes in the environment affect stored hippocampal maps. To this end, we changed path distances between object location during the path-change task, but not Euclidean distances. We expected that this would affect hippocampal representations in persons who are sensitive to egocentric features (response learners) more that it would in persons who are sensitive to allocentric features (place learners). Accordingly, we found that response learners updated representations in the left hippocampus based on the meaningful changes in path length that occurred in the path change task (Figure 6). Place learners, however, did not. Nevertheless, we did not find this effect when testing across all participants, highlighting an important role of navigation strategies.

First and foremost, these results show that a hippocampal map is not rigid, but can adapt flexibly to changes in the environment, which is crucial for effective navigation in a dynamic environment^1, 31^. Furthermore, these findings are consistent with the reported involvement of left hippocampus in sequential egocentric navigation^17, 18^, and support our earlier suggestion that navigational strategies might have an important influence on the nature of hippocampal cognitive maps. We speculate that the neural representations of response learners (egocentric navigators) might be more sensitive to changes in the environment that induce a change in egocentric behaviour such as changes in paths. Neural representations of place learners (allocentric navigators) might be less sensitive to such changes, as the allocentric map features, Euclidean distances, remain stable. We therefore speculate that navigational strategies might influence the sensitivity of hippocampal maps to environmental changes^38^. It might be that Euclidean (allocentric) changes impact allocentric navigators in a similar fashion as egocentric changes affect egocentric navigators. Interestingly, we found trend-level differences between place and response learners for distance representations in right hippocampus (after the object-location task). Together, these findings emphasise that it might be beneficial to investigate the influence of navigational strategies on hippocampal maps in future studies. Based on our results, we propose exploring whether inconsistencies in the literature might be due to differences between response and place learners.

Taken together, our results support the idea that hippocampal cognitive maps can adapt flexibly to changes in our environment. Furthermore, they highlight the important influence navigational strategies might have on hippocampal map representations.

## 5. Conclusion

We show that the hippocampus forms a map of our environment that represents, distinguishes and integrates Euclidean and path distance information. Importantly, we found evidence indicating that these maps can adapt flexibly to changes in the environment. The results of this study highlight the important role that navigational strategies and abilities might play in the formation and updating of hippocampal maps.

